# Increased context adjustment is associated with auditory sensitivities but not with autistic traits

**DOI:** 10.1101/2021.12.08.471716

**Authors:** R. Randeniya, J. B. Mattingley, M. I. Garrido

## Abstract

Bayesian models of autism suggest that disruptions in context-sensitive prediction error weighting may underpin sensory perceptual alterations, such as hypersensitivities. We used an auditory oddball paradigm with pure tones arising from high or low uncertainity contexts to determine whether autistic individuals display differences in context adjustment relative to neurotypicals. We did not find group differences in early prediction error responses indexed by mismatch negativity. However, the autism group had larger evoked responses to outliers, at 300ms latency suggesting a greater reorienting of attention to surprising sounds. A dimensional approach revealed a positive correlation between context-dependent prediction errors and auditory sensitivities, but not with autistic traits. These findings suggest that autism studies may benefit from accounting for sensory sensitivities in group comparisons.

**Lay Summary:** We find no differences in brain responses (EEG amplitudes) between autistic and neurotypical adults when listening to two contexts of tone patterns. However, we find a larger difference in the EEG amplitude when shifting between low to high uncertainity streams of tones, as sensory sensitivities (but not autistic traits) increase. These findings show that perceptual alterations maybe a function of sensory sensitivities, but not necessarily autistic traits. We suggest that future EEG studies in autism may benefit from accounting for sensory sensitivities.

## Introduction

Atypical sensory experiences are estimated to occur in over 90% of autistic children (Tomchek and Dunn, 2007) as well as in adults (Crane et al., 2009). Sensory disruptions are one of the first symptoms to appear in autism and have also been associated with core symptoms such as social communication deficits (Kern et al., 2007; Thye et al., 2018), which are positively correlated with autistic traits (Tavassoli et al., 2014b). Auditory hypersensitivities, one of the most commonly reported sensory anomalies, have been related to language developmental delays in autistic children (Jones et al., 2009; Eigsti and Fein, 2013) and have been shown to increase anxiety and limit participation in social activities (Stiegler and Davis, 2010). Understanding the mechanisms that give rise to such perceptual alterations will be useful in improving diagnostic tools and targeted interventions for autism.

“Bayesian Brain” perspectives of autism have given rise to models that attribute sensory perceptual dysfunctions in autism to an underlying disorder of precision (Lawson et al., 2014; Haker et al., 2016; Palmer et al., 2017). I.e., It is suggested that autistic perception is caused by altered reliability in observations or expectations (measured mathematically as precision, which is the inverse of variance of the representative Gaussian distribution) compared to neurotypicals. These theories suggest that sensory disruptions may arise either from forming poor models of the environment (Hypo-priors model; (Pellicano and Burr, 2012)) or due to sensory observations being too narrowly tuned (Precise likelihood model; (Brock, 2012)). Van De Cruys et al. (2014) suggest that the weighting of prediction error (i.e. the difference between our expectation of sensory information and the new sensory observation itself) is less flexibly adjusted in individuals on the autism spectrum, particularly across different contexts of uncertainty. This particular model, termed “Highly inflexible prediction errors in autism (HIPPEA)” is able to explain key symptoms of autism spectrum disorder (ASD) such as altered perceptual processing and resistance to change, as well as social deficits. Here we aimed to test this model and to investigate if prediction error updating between contexts is impaired in autism.

Mismatch negativity (MMN) is an ideal physiological marker for investigating sensory prediction errors (Garrido et al., 2009). The auditory MMN reflects a pre-attentive change detection in a pattern of stimuli and is particularly useful in that it can be studied without the confounding effects of attention (Näätänen et al., 2012) or task difficulty. There is currently limited literature on predictive processes in autistic adolescents and adults using the MMN, and of the studies that have been conducted the findings are mixed. Studies using classical oddball paradigms with pure tone frequency or duration deviants have demonstrated either no difference (Chien et al., 2018) or larger MMN (Lepistö et al., 2007) amplitudes in autistic individuals relative to controls. Other studies using more complex stimuli such as speech sounds (e.g., phonemes with or without affect) have also demonstrated either no difference in the mismatch response (Kasai et al., 2005) or reduced responses (Kujala et al., 2005) in ASD compared with neurotypicals. Two recent meta-analyses of both complex and pure tone findings showed no differences in MMN responses between autistic and neurotypical adults (Schwartz et al., 2018; Chen et al., 2020). Schwartz et al. (2018) add that their findings should be interpreted with caution because many studies included in their meta-analysis were underpowered; they also noted that there is a tendency for autistic adults to show larger MMN responses than their typically developing peers. Further, Schwartz et al. (2018) also found autistic (vs. neurotypical) groups to show greater differences in MMN to non-speech sounds compared with speech sounds. Very few studies, however, have investigated the flexibility of prediction error across different contexts in ASD. Goris et al. (2018), using an oddball paradigm, showed that global context modulated MMN in both ASD and control groups, but the effect was smaller in the autistic group compared with a neurotypical group. The study provides evidence that autistic individuals demonstrate impairment in context updating when the deviant occurs in more frequent vs. less frequent contexts.

In the present study our aim was to determine whether autistic individuals demonstrate prediction error adjustments to context, by evoking prediction errors (MMN) in contexts with low or high uncertainity. We used a stochastic oddball paradigm (Garrido et al., 2013) to compare: **a)** differential responses to standards and deviants (sensory prediction errors), which reflect an individual’s ability to learn about sensory context, their sensitivity to variability, and appropriate attribution of salience to odd but not common events, and **b)** MMN responses to the different levels of contextual precision that reflect an individual’s sensitivity to contextual uncertainty. The overall goal of the study was to provide evidence for or against models that describe perceptual disruptions as a disorder of prediction error weighting. We also took a dimensional approach in which we investigated both autistic traits and sensory sensitivities, the aim of which is to understand the relative contribution of sensory sensitivities to prediction error formation in autism.

## Methods

### Participants

Participants (N=59) between the ages of 18 - 35 years were recruited via Asperger’s Services Queensland, Autism Queensland and Mind and Hearts, The University of Queensland (UQ) online recruitment system, UQ newsletter, and online advertisements. Participants were recruited for the neurotypical (NT) group if they self-identified as having no diagnosis of neurodevelopmental (including an ASD) or psychiatric disorders and no current/history of medication acting on the nervous system. Participants with a reported diagnosis of an autism spectrum disorder only undertook an Autism Diagnostic Observation Schedule (ADOS; (Gotham, 2006; Hus and Lord, 2014)) interview with a trained clinical psychologist to confirm diagnosis. Participants who received a severity score below 3 (5 participants) were excluded from the Autism Spectrum (AS) Group. One participant did not complete the ADOS interview and thus was also excluded from the AS group. Thus, group comparisons were conducted with 23 AS and 23 age- and gender-matched NT participants. All participants (30 NT + 23 AS + 6 other) were included in the dimensional analysis of autistic and sensory sensitivity traits. This study was approved by the Human Research Ethics Committee of The University of Queensland (Approval No.: 2019000119).

### Procedure

#### Questionnaires

Self-report questionnaires included the Autism Quotient (AQ) questionnaire (Baron-Cohen et al., 2001) and the Sensory Processing Quotient (SPQ) Questionnaire (Tavassoli et al., 2014a), which were used to measure autistic traits and sensory sensitivities, respectively. It is important to note that the SPQ is a measure of thresholds and thus lower SPQ scores indicate more hypersensitivities. Participants also completed the Beck Anxiety Inventory (Beck et al., 1988) and Beck Depression Inventory (Beck et al., 1961).

#### Stochastic oddball paradigm

Participants underwent a stochastic frequency oddball paradigm (Garrido et al., 2013) and a simultaneous, visual 2-back task (Sweet, 2011) while undergoing an EEG recording. Participants listened to a stream of tones with log-frequencies sampled from two Gaussian distributions with equal means (500Hz) and different standard deviations (*narrow*: *σ*_*n*_ = .5 octaves; *broad*: *σ*_*b*_ = 1.5 octaves); see Figure 1. Thus, participants heard frequencies drawn from two contexts of either high uncertainty (200Hz to 4000Hz) or low uncertainty (200Hz to 2000Hz). The mean values of both Gaussian distributions (500Hz) were the ‘standard’ tones and constituted 10% of all the tones (i.e., 200 tones). 10% of the tones which were outliers to the Gaussian distributions were defined as ‘deviant’ tones, which were played at 2000Hz. All tones had a duration of 50ms with 10ms smooth rise and fall periods and inter-stimulus intervals of 500ms. Standard and deviant tones occurred randomly in the stream of tones. Participants were instructed to disregard the tones and to focus instead on the visual 2-back task, in which they had to press a button on a keyboard if any letter on the computer screen repeated after 2 letters (visual 2-back task). This experimental component lasted for approximately 30 minutes and was divided into 4 blocks (2 narrow and 2 broad) with short breaks in between each block. The *narrow* and *broad* distribution blocks were counter-balanced across participants.

**Figure 1:**
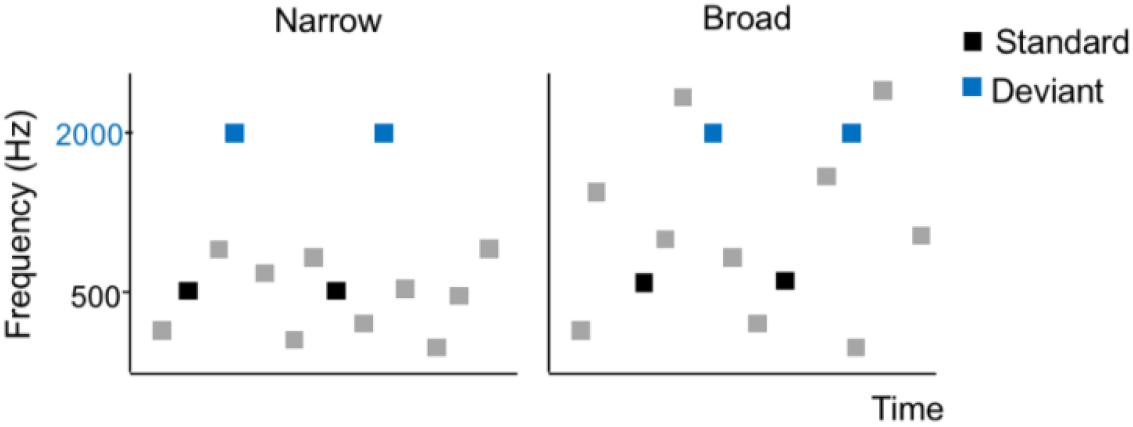
Stochastic Oddball paradigm. Participants listened to a stream of 500ms tones drawn from either a Narrow (left) or Broad (right) distribution of frequencies. 500Hz (black) tones were standards and 2000Hz (blue) tones were deviants.

#### EEG Data Acquisition and Processing

Throughout the auditory oddball experiment, an electroencephalogram (EEG) was recorded using a 64 electrode-cap with Bio Semi Act iView system at a sampling rate of 1024Hz and the following specifications: 417Hz bandwidth (3dB) and 18dB/octave roll-off. Further electrodes were placed on the outer canthi of both eyes, as well as below and above the left eye to measure eye movement. Triggers were marked in the EEG data at the onset of each tone. Raw EEG data was first filtered using a band pass filter between 0.5Hz – 40Hz. Data were segmented into 500ms epochs (including 100ms prestimulus baseline and 400ms from the onset of the stimulus). Epochs containing artefacts exceeding ±50 V were excluded. Trials were next averaged together by condition (Narrow Standard, Narrow Deviant, Broad Standard and Broad Deviant) and baseline corrected using a prestimulus interval of 100ms. Data were pre-processed and analysed using SPM 12 (Wellcome Trust Centre for Neuroimaging, London; http://www.fil.ion.ucl.ac.uk/spm/) in MATLAB version R2018b.

##### Whole scalp analysis

We undertook a whole scalp analysis using SPM 12. Averaged epochs for each condition and participant were converted into 3D spatiotemporal images. The spatiotemporal image volumes were modelled with a general linear model of 2×2×2 ANOVA design, with the factors of Group (AS vs NT), Context (Narrow vs Broad) and Surprise (Standards vs Deviants).

##### Single Channel Analysis (Fz) electrode

To enable comparison with previous literature (Schwartz et al., 2018) we also undertook a single channel analysis by measuring amplitudes for each condition at the Fz electrode. MMN was measured as the mean amplitude(μV) in difference of deviants and standards between 125ms and 175ms.

We further conducted an exploratory analysis, given that we observed differences in a negative deflection in the prediction error waveform (Deviants minus Standards) at 300ms after stimulus onset. We term this component N300 as it is a negative deflection at 300ms. This N300 component was measured as the mean amplitude(μV) in difference of deviants and standards between 300ms and 380ms.

For group analysis we conducted 2×2×2 ANOVA with factors of Group (AS vs. NT), Context (Narrow vs. Broad) and Surprise (Deviants vs. Surprise). For trait analysis we first conducted a regression analysis with delta-MMN or delta-N300 as the outcome variable and AQ scores, SPQ auditory scores, Group membership (NT/AS/Other), and medication use as predictor variables. We included both AQ and SPQ auditory scores in the same model as they were not correlated with each other. Where a significant regression was identified we conducted Pearson’s correlation analysis to understand the relative contributions of standards and deviants on the predictor. All statistical analyses were run in MATLAB R2018b and figures are presented using ggplot2 (Wickham, 2016) in RStudio.

## RESULTS

### Participants

ADOS interview confirmed 23 participants to be on the autism spectrum, and these participants were included in the AS group (Age *M* = 24.35 , *SD* = 6.08; 12 females, 10 males, 1 intersex). The NT group consisted of 23 age- and gender-matched controls (Age *M* = 24.04, *SD* =6.06; 12 females, 11 males); see Table 1 and Figure 2 for demographic details. The AS group showed more Anxiety [*t* = 1.697 , *p* = 0.097, *BF* =1.306] and higher Depression Scores [*t* = 2.636 , *p* = 0.012, *BF* = 0.248] as well as AQ scores [*t* = 6.861, *p* = 1.829×10^−8^, *BF* = 2 x 10^−6^]. The AS group showed no differences in auditory hypersensitivities (i.e., SPQ auditory subscale score) compared with the NT group [*t* = − 0.080, *p* = 0.936, *BF* = 4.560]. Eleven participants in the AS group reported using antidepressants or anti-anxiety medication (e.g., Escitalopram, Venlafaxine, Setraline, Amitriptyline or similar drugs of serotonin reuptake inhibitor class) and 6 participants reported ADHD medication (i.e., Methylphenidate class).

**Table 1:**
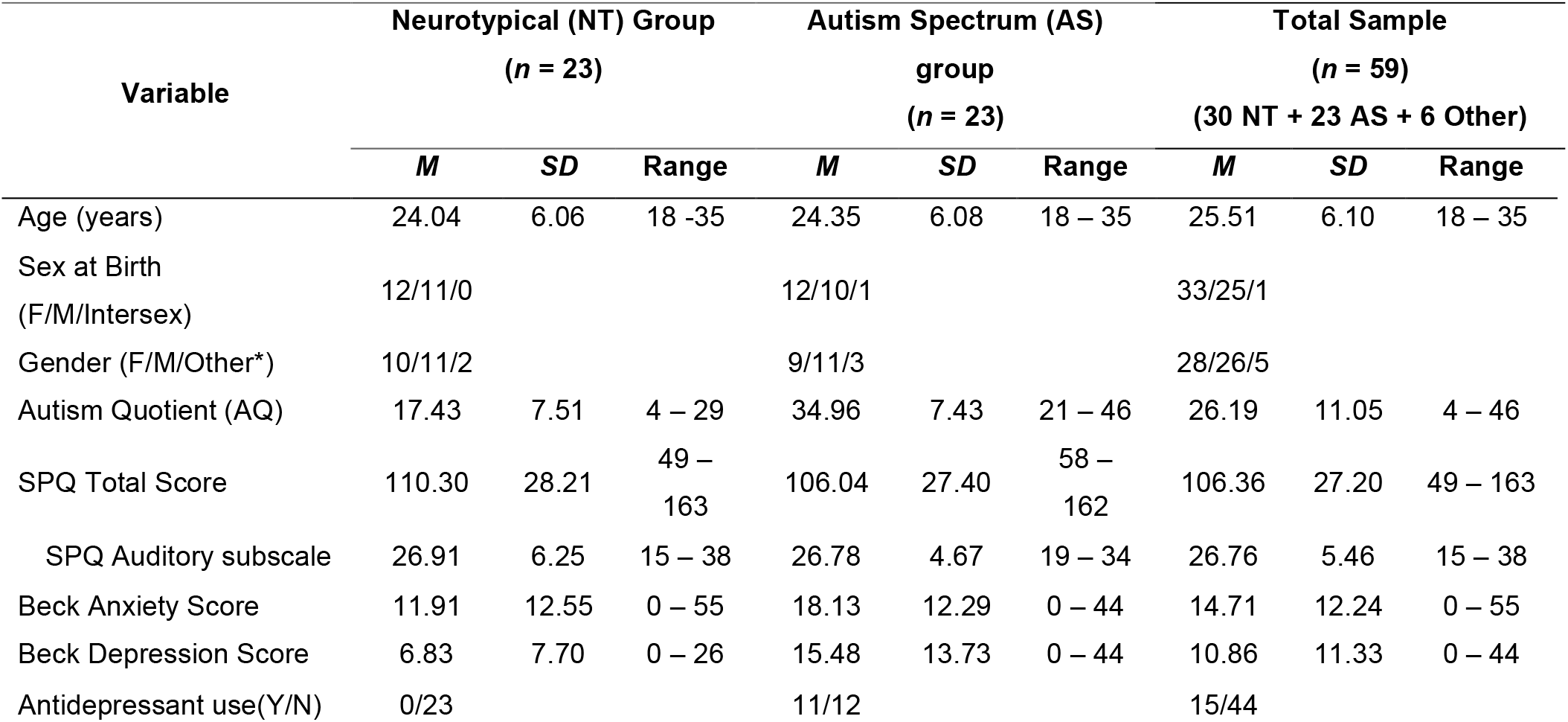
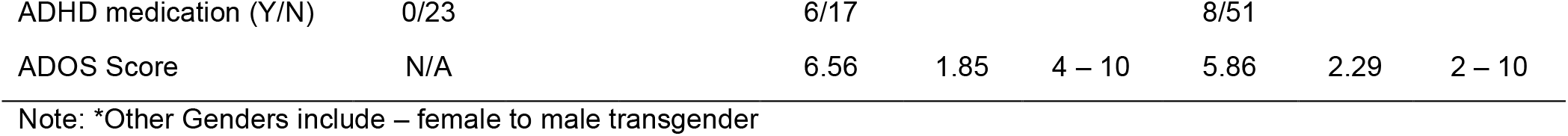
Demographic Details.

**Figure 2:**
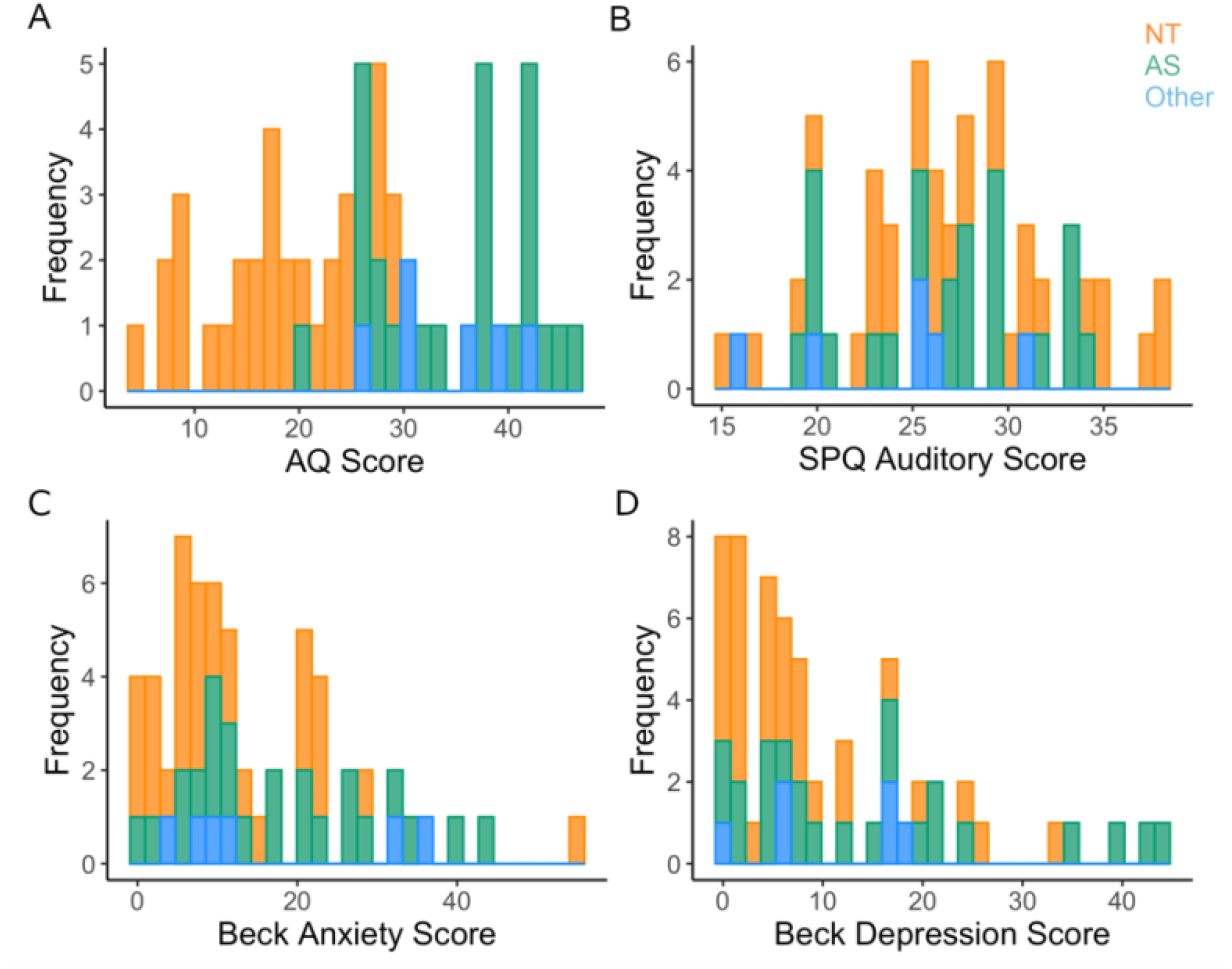
Psychometric profile of participants. A) Autism Quotient (AQ) B) Sensory Processing Quotient - auditory subscale score C) Beck Anxiety Inventory Score d) Beck Depression Inventory for neurotypicals (NT; orange), confirmed autistic (AS; green) and participants who identified as having received a diagnosis of an autism-spectrum disorder but which could not be confirmed during interview with a psychologist (Other; blue).

In the full sample (N = 59 ; i.e., NT = 30, AS = 23, Other = 6; Figure 2), AQ scores were significantly correlated with Anxiety [*r* =0.360, *p* = 0.014 , *BF* = 0.432] and Depression [*r* =0.422, *p* = 0.003, *BF* = 0.125] but not with SPQ auditory scores [*r* =−0.053, *p* = 0.726, *BF* = 8.163].

#### Whole Scalp Results

A 2×2×2 ANOVA of EEG activity revealed significant clusters at the whole-scalp *p*_FWE_ <0.05 threshold for the main effect of *Surprise* arising over fronto-central channels at 165ms [cluster size k_E_ = 10,825, *p*_cluster-FWE_ = <0.001, *F* = 45.38, *Z* = 6.24], parieto-occipital channels at 149ms [k_E_ = 4006 , *p*_cluster-FWE_ = <0.001, *F* = 40.65, *Z* = 5.93] and at occipital channels at 366ms [k_E_ = 513 , *p*_cluster-FWE_ = 0.004, *F* = 25.12, *Z* = 4.70] and at 346ms [k_E_ = 141 , *p*_cluster-FWE_ = 0.017, *F* = 19.95, *Z* = 4.19]. We also observed significant clusters in *Context*Surprise* interactions over fronto-central channels at 162ms [k_E_ = 1596, *p*_cluster-FWE_ = <0.001, *F* = 25.30, *Z* = 4.72] and 241ms [k_E_ = 463, *p*_cluster-FWE_ = 0.005, *F* = 23.80, *Z* = 4.58] and parieto-occipital channels at 170ms [k_E_ = 137, *p*_cluster-FWE_ = 0.005, *F* = 21.25, *Z* = 4.32]. This confirmed that the paradigm elicited context-specific prediction error responses, as expected.

We found no significant clusters for *Group*Surprise* or *Group*Context*Surprise* interactions at the whole-scalp *p*_FWE_ <0.05 or *p*_uncor_ < 0.001 levels, indicating no evidence for group differences in MMN across contexts.

#### Single Channel (Fz) analysis

##### No group differences in MMN but differences in N300

A 2×2×2 ANOVA of mean amplitudes at the Fz channel in the MMN window (see methods) revealed no significant *Group*\**Context*\**Surprise* [Effect Size η_p_^2^ = 0.062, *p* = 0.096, *F* = 2.892], *Group*\**Surprise* [η_p_^2^ = 0.022, *p* = 0.320, *F* = 1.008], or *Group*\**Context* [η_p_^2^ = 0.020, *p* = 0.348, *F* = 0.899] interactions. Thus, there was no evidence for group differences in MMN, and no differences in context adjustment (delta-MMN). See Figure 3 prediction error waveforms at Fz.

**Figure 3:**
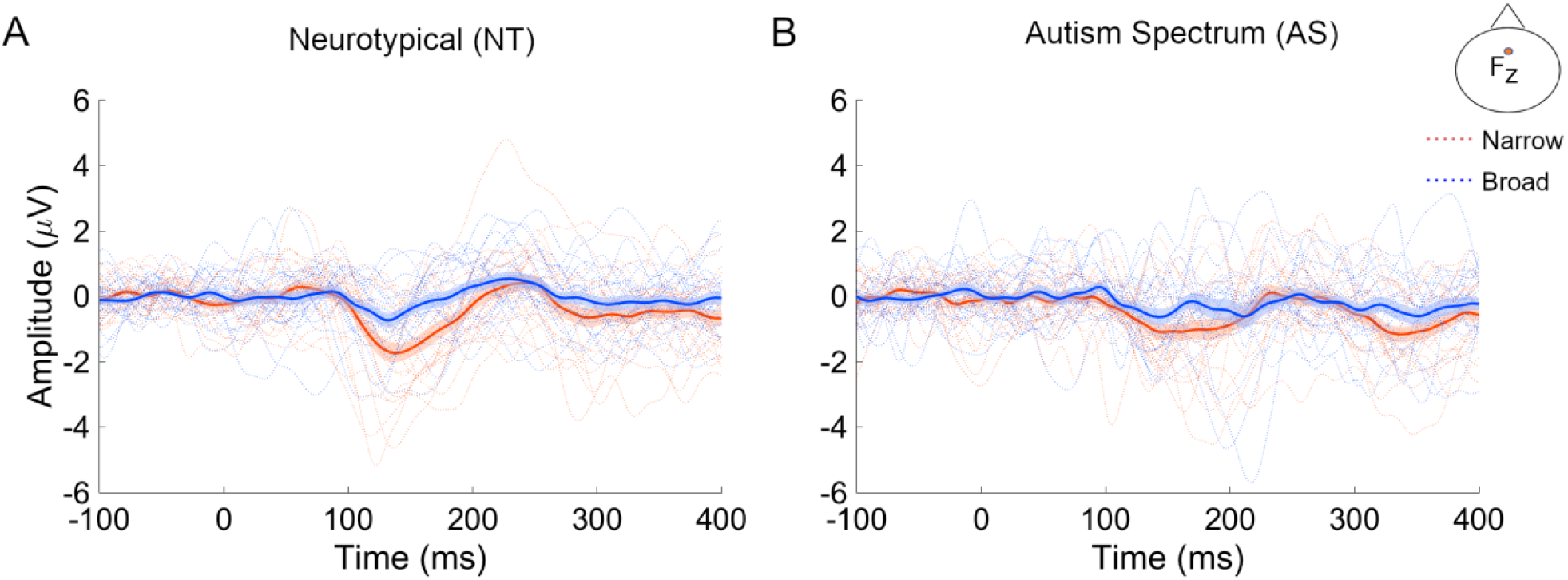
Prediction error waveforms at Fz electrode. Mismatch response (difference in Deviants > Standards) for A) matched Neurotypical (NT) and B) Autistic (AS) groups for narrow (red) and broad (wide) oddball contexts. Location of Fz electro3de on scalp is shown at top right.

We also investigated amplitudes at a N300 window. We observed a medium effect for *Group*\**Surprise* [η_p_^2^ = 0.098, *p* = 0.034, *F* = 4.776] but no evidence for a *Group*Context* [η_p_^2^ = 0.017, *p* = 0.390, *F* = 0.755] or *Group*\**Context*\**Surprise* [η_p_^2^ = 0.018, *p* = 0.369, *F* = 0.823] interaction. Thus we observed no differences in delta-N300 but there were significant differences in the overall surprise effect between groups, across conditions. Post-hoc analysis of this *Group*\**Surprise* interaction revealed the AS group (vs. NT) had a more negative across all conditions compared to NT group [Mean difference = 1.1107, *t* = 2.185].

##### Auditory sensitivities but not autistic traits are correlated with delta-MMN

A regression analysis with AQ and SPQ scores as predictors of delta-MMN amplitude [*F* = 5.823, *p* = 0.005] revealed auditory SPQ scores were a significant predictor of delta-MMN amplitudes [*Beta* = 0.419, *p* = 0.008] even when adjusted for AQ Scores, Anxiety, Depression, medication use and group membership (i.e., NT/AS/Other); See Table 2. This indicates that as auditory sensitivities increased (lower SPQ scores) participants showed a larger difference in MMN between contexts. We further investigated if this delta-MMN relationship with auditory sensitivities could be attributed to the standards or deviants. We found that there was no correlation with the auditory SPQ scores and delta-Standards [*r* = −0.192, *p* = 0.145, *BF* = 3.403], indicating that model formation was not modulated by auditory sensitivities. However, we observed a weak positive correlation with the delta-deviants [*r* = 0.262, *p* = 0.045, *BF* = 1.335] (Figure 4) suggesting context-dependent surprise responses increase with increasing sensory sensitivity (decreasing SPQ scores). We also found no relationship between ADOS severity scores and delta-MMN [*r* = 0.299, *p* = 0.122 , *BF* = 2.093] or delta-N300 [*r* = −0.169 , *p* = 0.391 , *BF* = 4.753].

**Table 2:** Linear regression model predicting delta-MMN amplitude.

| Predictor Variables | Beta | SE | *p* | 95% confidence |  |
| --- | --- | --- | --- | --- | --- |
|  |  |  |  | interval |  |
|  |  |  |  | Lower | Upper |
| Model 1 |  |  |  |  |  |
| AQ Score | 0.231 | 0.018 | 0.143 | -0.010 | 0.061 |
| SPQ Auditory Score | 0.383 | 0.030 | 0.003 | 0.030 | 0.150 |
| Model 2 |  |  |  |  |  |
| AQ Score | 0.389 | 0.026 | 0.108 | -0.008 | 0.093 |
| SPQ Auditory Score | 0.419 | 0.034 | 0.008 | 0.034 | 0.172 |
| Beck Anxiety | 0.080 | 0.022 | 0.693 | -0.043 | 0.045 |
| Beck Depression | -0.111 | 0.019 | 0.449 | -0.043 | 0.033 |
| Anx/Dep Medication | -0.095 | 0.467 | 0.554 | -1.205 | 0.593 |
| ADHD Medication | -0.235 | 0.725 | 0.208 | -2.516 | 0.336 |
| Group (AS/NT/Other) | -0.064 | 0.362 | 0.742 | -0.862 | 0.590 |

Outcome variable: delta-MMN

**Figure 4:**
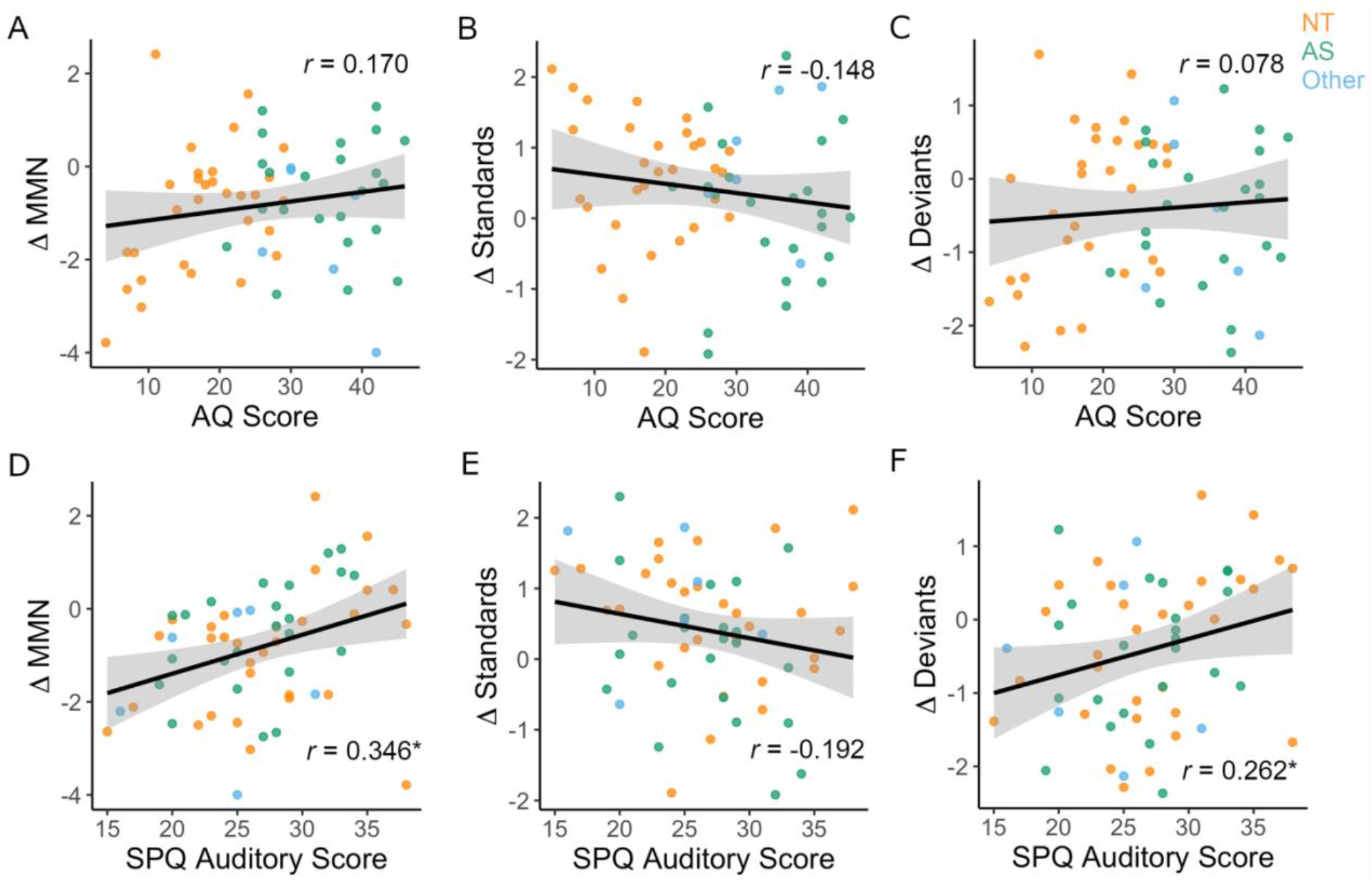
Trait Analysis. Correlations between autism quotient (top, panels A, B, C) and SPQ auditory subscale scores (bottom, panels D, E, F) with mean amplitude (μV) in the mismatch negativity time window. Differences between broad and narrow conditions are shown for delta-MMN, delta-standards and delta-deviant amplitudes. Neurotypical (NT; orange), autistic (AS; green) and unconfirmed autistic (Other; blue).

We also conducted a regression analysis (See methods) with AQ Score and SPQ Score as predictors of delta-N300 amplitude (see Table 3). Only Group membership (AS/NT/Other) was a significant predictor of delta-N300 [*Beta* = 0.489, *p* = 0.019].

**Table 3:** Linear regression model predicting delta-N300 amplitude.

| Predictor Variables | Beta | SE | $p$ | 95% confidence interval |
| --- | --- | --- | --- | --- |

|  |  |  |  | Lower | Upper |
| --- | --- | --- | --- | --- | --- |
| **Model 1** |  |  |  |  |  |
| AQ Score | 0.170 | 0.017 | 0.183 | -0.008 | 0.059 |
| SPQ Auditory Score | -0.114 | 0.037 | 0.409 | -0.094 | 0.052 |
| **Model 2** |  |  |  |  |  |
| AQ Score | 0.043 | 0.027 | 0.841 | -0.044 | 0.063 |
| SPQ Auditory Score | -0.037 | 0.035 | 0.772 | -0.073 | 0.062 |
| Beck Anxiety | -0.07 | 0.021 | 0.638 | -0.058 | 0.032 |
| Beck Depression | -0.077 | 0.018 | 0.564 | -0.046 | 0.027 |
| Anx/Dep Medication | -0.201 | 0.674 | 0.322 | -2.152 | 0.547 |
| ADHD Medication | -0.172 | 0.848 | 0.408 | -2.252 | 0.987 |
| Group (AS/NT/Other) | 0.489 | 0.468 | 0.019 | 0.068 | 1.924 |

Outcome variable: delta-N300

##### No group differences or dimensional associations in variability of amplitude of standards

We also investigated the variability of mean amplitudes of standard trials in the MMN time window across trials. We used the first 50 trials to assess if there was variability in standards which would indicate differences in the initial learning of a pattern (i.e., forming a prior). We found no difference between groups in either the *Narrow Standards* [Mean difference = 15.413, *SD* = 17.723, *t* = 0.870, *p* = 0.389, *BF* = 3.269] or in *Broad Standards* [Mean difference= −80.9837, *SD* = 104.837, *t* = −0.772, *p* = 0.444, *BF* = 3.508] conditions. We also found no relationship between AQ Scores and variability in the *Narrow Standards* [*r* = 0.091, *p* = 0.495 , *BF* = 8.768] and *Broad Standards* [*r* = −0.119, *p* = 0.367, *BF* = 3.768] conditions, nor did we observe differences with SPQ auditory scores and variability in the *Narrow Standards* [*r* = −0.217, *p* = 0.099, *BF* = 2.533] and *Broad Standards* [*r* = − 0.130, *p* = 0.325, *BF* = 6.049] conditions. We also found no differences with ADOS scores and the variance of standards in the *Narrow Standards* [*r* = 0.033, *p* = 0.867, *BF* = 6.759] and *Broad Standards* [*r* = 0.092, *p* = 0.647, *BF* = 6.148] conditions.

## Discussion

In this study we aimed to understand if prediction error generation is disrupted in autism and also if autistic individuals display anomalies in context adjustment to uncertainty relative to NT controls. We observed no group differences in the classical MMN within conditions, but found larger N300 responses for the AS than the NT group. The MMN findings are in line with recent meta-analysis findings (Schwartz et al., 2018; Chen et al., 2020). We also investigated the flexibility of prediction errors across contexts with varying uncertainty (low vs. high precision contexts). Again, we found no differences between AS and NT groups in adjusting prediction errors between contexts in the MMN window, which is in contradiction to (Goris et al., 2018), who find a reduction in MMN amplitude. These findings suggest that both model forming and context adjustment may be intact in autism. Further, when taking a dimensional approach, we found that autism traits did not predict context adjustment (delta-MMN). These contrasting findings from a categorical and dimensional approach may be due to our two groups (AS and NT) not differring in their SPQ auditory scores. These findings are important in that they may explain the contrasting findings of low vs. high MMN amplitudes in autistic individuals compared with neurotypicals. Furthermore, these findings highlight the importance of characterizing sensory symptoms in autistic and neurotypical individuals, and accounting for these differences in group comparisons. Further, context adjustment associations with SPQ scores provide evidence for the prediction error weighting hypothesis, but do not demonstrate inflexibility to context, thus providing partial evidence against the HIPPEA (Van de Cruys et al., 2014) model of hypersensitivities.

We also provide evidence for intact priors in autism in that standards across contexts showed no differences, and there were no differences in their variability. Thus we provide strong evidence against the hypo-priors model, which suggests that autistic perception is characterized by forming less precise models of the world (Pellicano and Burr, 2012). While this has not been shown previously using MMN specifically, evidence for intact priors in autism have been demonstrated in visual (Pell et al., 2016; Croydon et al., 2017; Karvelis et al., 2018) and tactile sensory (Cannon et al., 2021) modalities. Similar to Butler et al. (2017), our findings reveal no variability differences in standards on a trial-by-trial basis, which provide evidence against a sensory unreliability hypothesis in autism.

We also conducted an exploratory analysis in an N300 window given that we observed larger negative deflections in the prediction error waveforms of the autistic group. This component is not typically observed between 300-400ms, where usually a positive deflection in the prediction error waveform (termed P300) would be apparent. The P300/P3a has been shown to have no differences in autistic adolescents and adults compared with neurotypicals, and P3a amplitudes have been shown to be correlated with sensory avoidance (Chien et al., 2018). This N300 component could also reflect an early reorienting negativity (RON), which is typically observed at 400 – 600ms post stimulus onset (Berti, 2008), which is thought to reflect attention orienting but has been not been widely studied in autism. Nonetheless, we observed significant group differences in overall surprise response at 300ms, where the AS group showed larger N300 relative to the NT group, indicating that our AS group showed a larger orienting response to deviant tones. However, we did not observe any relationships between autistic traits/sensory sensitivities and delta-N300, suggesting that the N300 may be capturing an aspect specific to the diagnosed autistic group.

There are a number of caveats to consider in relating our findings to hypotheses arising from Bayesian implementations of sensory learning in autism. First, mismatch negativity amplitudes have been shown to be influenced by factors such as medication. More than 50% of our AS group participants reported antidepressant use. Selective serotonin reuptake inhibitor (SSRI) class drugs, such as Escitalopram have been shown to increase MMN amplitudes in healthy individuals (Oranje et al., 2008; Wienberg et al., 2010). A smaller number of our AS group participants also reported use of attention modulating drugs of the Methylphenidate class, such as Ritalin. Methylphenidates have been shown to reduce group differences in event-related potential indices in children with ADHD (Ozdag et al., 2004; Lawrence et al., 2005). Thus, further study of prediction errors using MMN in larger samples with drug-naïve autistic participants will be important for understanding the relative contributions of medication to sensory processing. Further, MMN amplitudes have also been shown to decline with age (Cheng et al., 2013b; Cheng et al., 2013a). This limits comparability of our findings in an adult sample with MMN findings in children. A global theory of prediction error adjustment relevant to autism would ideally be validated across age groups.

In summary, our study demonstrates the importance of undertaking a dimensional approach, specifically taking sensory sensitivities into account when investigating uncertainty under different contexts. Schwartz et al. (2018) pointed out that within-group variability of MMN anomalies may serve as a better avenue for study than between-group analysis. Our findings also provide evidence for predictive processes above and beyond simple habituation deficits which may describe sensory sensitivities and underpin the importance of studying predictive processes in autism under varying contexts.

## Acknowledgements

We thank Radhika Tanksale for conducting ADOS assessments, and our participants for their valuable time. This work was supported by the Australian Research Council Centre of Excellence for Integrative Brain Function (ARC Centre Grant CE140100007; MIG and JBM). RR was supported by The University of Queensland Research Training Programme. MIG was supported by a University of Queensland Fellowship (2016000071).

